# Thalamic encoding of lexical status is lateralized during reading aloud

**DOI:** 10.1101/2020.07.30.229898

**Authors:** Dengyu Wang, Witold J. Lipski, Alan Bush, Anna Chrabaszcz, Christina Dastolfo-Hromack, Michael W. Dickey, Julie A. Fiez, R. Mark Richardson

## Abstract

To explore whether the thalamus participates in lexical status encoding, local field potentials were recorded in patients undergoing deep brain stimulation lead implantation while they read aloud single-syllable words and nonwords. Bilateral decreases in thalamic beta (12-30Hz) activity were preferentially locked to stimulus presentation, and these decreases were greater when nonwords were read. Increased broadband gamma (70-150Hz) activity also was locked preferentially to speech onset bilaterally, but greater nonword-related increases in this activity were observed only on the left, demonstrating lateralization of thalamic gamma selectivity for lexical status. In addition, this lexical status effect was strongest in more anterior thalamic locations, regions which are more likely to receive pallidal than cerebellar afferents. These results provide evidence from intracranial thalamic recordings for the lateralization and topography of subcortical lexical status processing.

## Introduction

Reading words aloud depends on our ability to transform information about letter combinations into plans for producing speech sounds. Determining whether a group of letters represents a word is one important component of this process. While word reading can be supported by processes that permit mapping entire word forms to lexical representations, nonword reading depends upon sublexical processes that map spelling and sound (Coltheart et al., 2001). Functional neuroimaging has allowed for increasingly detailed study of the cortical regions that participate in these phonological processes, for instance demonstrating that a region of the inferior frontal gyrus (Brodmann’s areas 44 and 45) is significantly more active for nonword reading than for word reading (Fiez et al., 1999; Hagoort et al., 1999; Heim et al., 2005; Herbster et al., 1997). The role of subcortical regions in spoken word production remains elusive, however, due to the low resolution of neuroimaging techniques for measuring subcortical activity. Resolving this knowledge gap is important, given that cortical activity is modulated by thalamic outflow through basal ganglia-thalamo-cortical and cerebello-thalamo-cortical circuits (Behrens et al., 2003; Bosch-Bouju et al., 2013; Hwang et al., 2017; Zhang et al., 2010). Despite the cortico-centric focus of most experimental work and accompanying models, there is increasing recognition of the role of subcortical processes in language production (Llano, 2015).

Neurosurgical procedures involving invasive recording and stimulation in epilepsy patients undergoing electrode implantation for seizure mapping traditionally have provided the only direct means to test hypotheses related to cortical function during reading (Juphard et al., 2011). Epilepsy surgery, however, rarely provides access to the thalamus and basal ganglia. Movement disorders surgery, on the other hand, routinely provides direct access to the thalamus and basal ganglia in awake patients. Regionalization of language function within the left thalamus was established in surgery for movement disorders in the 1970s, in seminal studies by Ojemann and colleagues that employed electrical stimulation principles borrowed from traditional cortical language mapping protocols (Johnson & Ojemann, 2000). More recently, event-related potential recordings in patients undergoing deep brain stimulation (DBS) have suggested that thalamic structures are engaged in the analysis of syntactic, semantic and lexical information during acoustically presented language tasks (Tiedt et al., 2017; Wahl et al., 2008).

We recently developed a protocol to study subcortical activity during single-syllable word/nonword reading, in patients undergoing DBS lead implantation (Chrabaszcz et al., 2019; Lipski et al., 2018). Here, we explore whether the thalamus participates in the encoding of lexical status, by recording local field potentials (LFP) in patients undergoing DBS lead implantation for essential tremor targeting the ventral intermediate nucleus of the thalamus (Vim, which corresponds to the ventral portion of the ventral lateral posterior nucleus (VLp) (Macchi & Jones, 1997)). Subjects performed a reading aloud task where they were asked to read aloud single-syllable words or nonwords that appeared on a computer screen (Figure 1). We assessed thalamic participation in lexical status encoding by comparing task-related neural responses when participants spoke nonwords vs. words.

**Figure 1:**
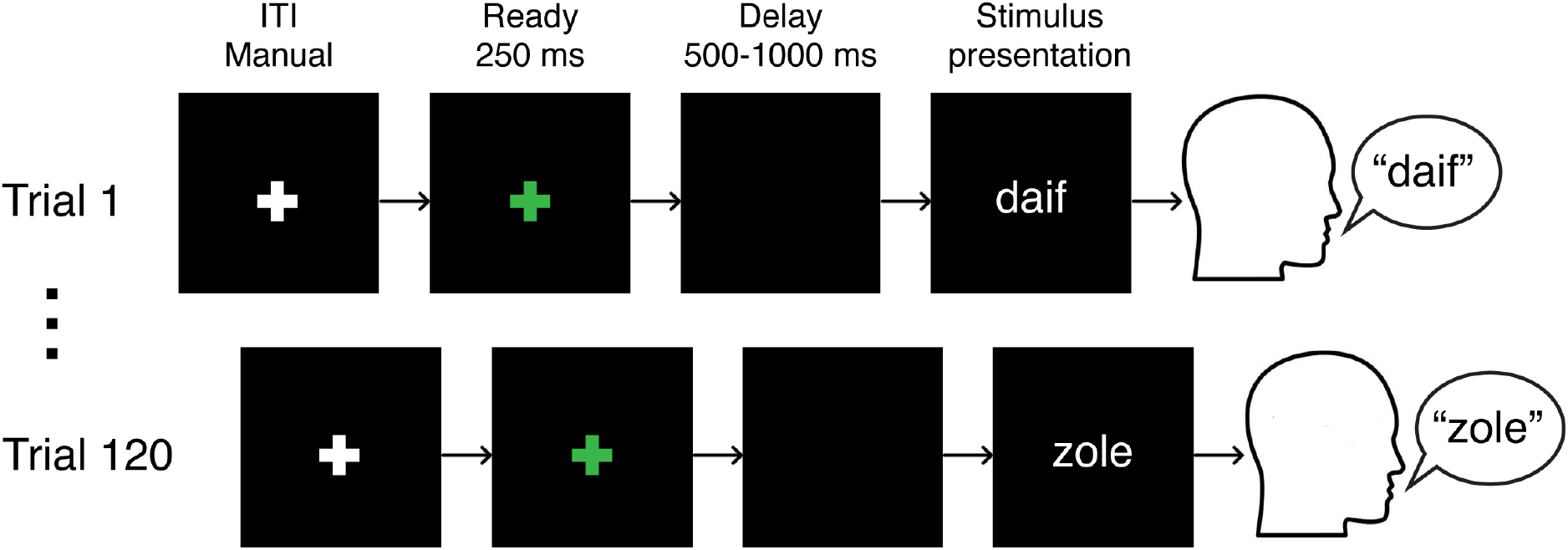
Schematic of experiment. ITI: inter-trial interval.

## Results

Eleven subjects read aloud single-syllable words alternating with nonwords, intraoperatively during implantation of DBS leads targeting the Vim nucleus of the thalamus. LFP recordings in four subjects were obtained unilaterally (3 left, 1 right) during microelectrode mapping, where each of these subjects performed up to four sessions of 120 trials. Recordings from seven subjects were obtained from DBS lead contacts, where each of these subjects performed two sessions: in the first session unilateral recordings were made from the left thalamus, and in the second session bilateral recordings were preformed simultaneously. A total of 117 recordings (data recorded in one location in one session) from 89 recording sites pooled across subjects were collected. Recording locations were determined in MNI (Montreal Neurological Institute) space (Figure 2) and comprised locations within or bordering (within 1mm) the ventral anterior nucleus (VA) and the ventral lateral anterior nucleus (VLa) (38/89), or the VLp (51/89).

**Figure 2:**
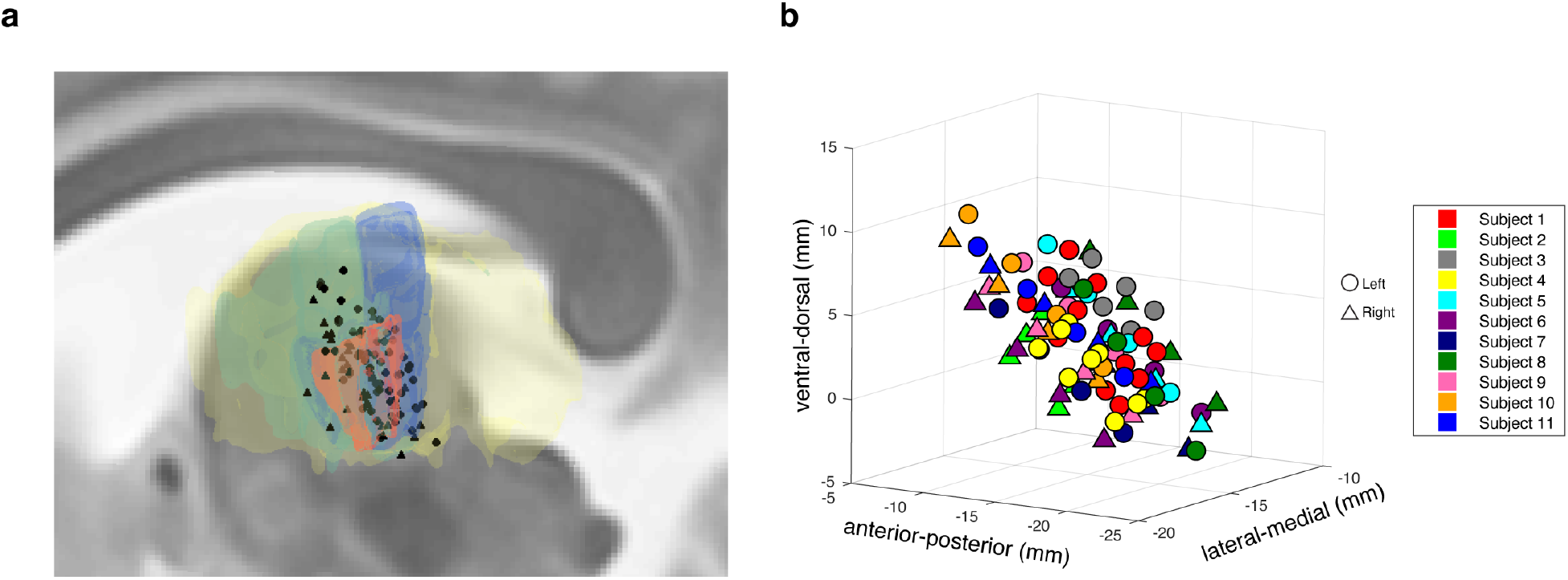
Localization and MNI-transformation of recording sites. **a** Sagittal view of the recording locations of all the subjects relative to the thalamus (yellow), VA and VLa (green), VLp (blue), and Vim (orange), superimposed on a T2-weighted image. **b** A plot of all the recording sites in MNI space with their MNI-defined coordinates. Recording contacts of different subjects are color-coded. In both **a** and **b**, right hemisphere recording locations are flipped to the left hemisphere: left side contacts are marked with circles, and right side contacts are marked with triangles.

### Behavioral performance

Across all subjects, the mean speech production latency (interval between stimulus presentation and onset of speech) was 1.07 ±0.29s, and the mean duration of speech was 0.70 ±0.09s. Nonword production latency (1.23 ±0.37s) was significantly longer than word production latency (0.98 ±0.24s) across subjects (two-tailed paired t-test, t(10)=4.47, *P*=0.0012; Figure 3a). Nonword production duration across subjects (0.68 ±0.07s) was not significantly different from word production duration (0.72 ±0.13s; two-tailed paired t-test, t(10)=1.99, *P*=0.074; Figure 3b).

**Figure 3:**
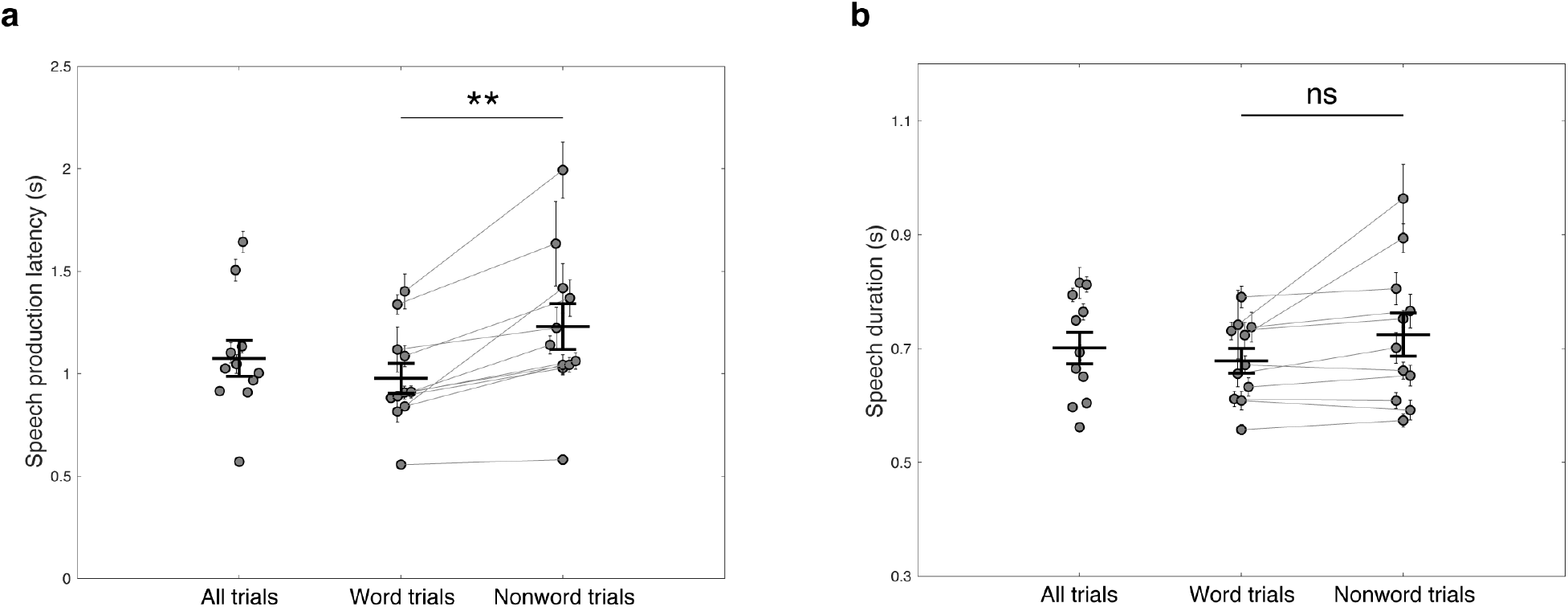
Behavioral outcomes. **a** Mean and SEM of speech production latency for each subject across all trials (first column), word trials (second column), or nonword trials (third column), superimposed with the mean and SEM of speech production latency across subjects. **b** Mean and SEM of speech duration for each subject across all trials (first column), word trials (second column), or nonword trials (third column), superimposed with the mean and SEM of speech duration across subjects. Two-tailed paired t-test, \*\**P*<0.01, ns: not significant.

### Thalamic neural activity is modulated during reading aloud

Thalamic LFP activity exhibited significant time-frequency modulation during the reading aloud task (Figure 4). Compared to baseline (a period of 1000ms preceding stimulus presentation), there was a significant decrease in spectral power in the beta frequency band (12-30Hz) that occurred at stimulus presentation and lasted until the end of speech (−1.08-0.62s relative to speech onset, two-tailed Wilcoxon signed-rank test, n=117, *P*<0.05, Bonferroni corrected). In contrast, a significant increase in broadband gamma (70-150Hz) activity occurred shortly before the onset of speech and persisted throughout the utterance (−0.15-0.59s relative to speech onset, two-tailed Wilcoxon signed-rank test, n=117, *P*<0.05, Bonferroni corrected). Average z-scored task-related beta and broadband gamma response amplitudes of each trial were then calculated over the respective significant time windows for all the recordings. As a result, 66/117 (56%) of the recordings showed significant beta activity decreases during the task compared to baseline (one-tailed one-sample t-test, *P*<0.05, Bonferroni corrected), and significant task-related broadband gamma activity increases were observed in 91/117 (78%) of the recordings (one-tailed one-sample t-test, *P*<0.05, Bonferroni corrected).

**Figure 4:**
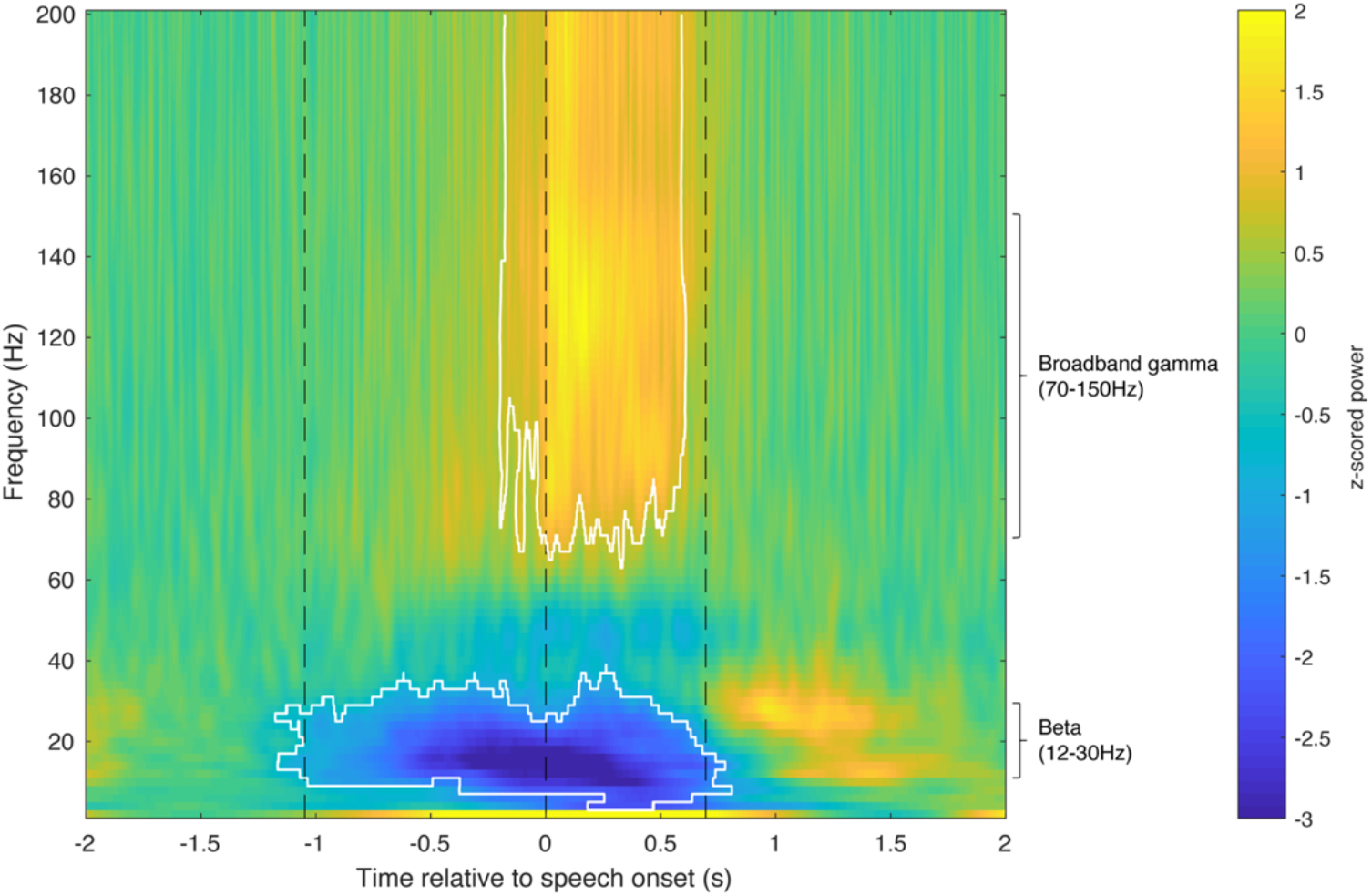
Thalamic neural activity shows task-related modulations. A spectrogram of thalamic neural activity during the reading aloud task, averaged across all trials and all recordings. Trials are aligned to speech onset. Significant changes compared to baseline are marked in white contours (−1.08-0.62s for beta activity, and −0.15-0.59s for broadband gamma activity; Wilcoxon signed-rank test, n=117, *P*<0.05, Bonferroni corrected). Average time points of stimulus presentation, speech onset and offset are marked with black dashed lines.

### Beta and broadband gamma responses differ in timing properties

The average interval between stimulus presentation and onset of a significant change from baseline in the spectral power of a particular frequency band, i.e. the mean band response latency, was shorter for significant beta decrease responses than for significant broadband gamma increase responses (0.79 ±0.18s vs. 0.99 ±0.18s, two-tailed two-sample t-test, t(155)=-6.97, *P*<10^−5^). The mean band response onset to speech onset interval was also greater for significant beta decrease responses than for significant broadband gamma increase responses (0.31 ±0.15s vs. 0.13 ±0.14s, two-tailed two-sample t-test, t(155)=7.67, *P*<10^−5^). To characterize the temporal properties of these responses, we examined their trial-to-trial relationships to stimulus presentation versus speech onset (Figure 5a-d). Of the 66 recordings that showed significant task-related beta power decreases, 43 (65.2%) had beta responses time-locked to stimulus presentation, whereas only 19 (28.8%) had beta responses time-locked to speech onset (Figure 5e). In contrast, the majority (70/91, 77.0%) of significant broadband gamma power increases were time-locked to speech onset, with a minority (19/91, 21.0%) time-locked to stimulus presentation (Figure 5f). These relationships were dissociated (χ^2^ test, α=0.05; Supplementary Table 1), with beta decreases more likely to be stimulus-locked (χ^2^(1)=31.4, *P*<10^−5^) and broadband gamma increases more likely to be speech onset-locked (χ^2^(1)=36.1, *P*<10^−5^). These temporal correlations did not differ between recording sides (χ^2^ test, α=0.05; Supplementary Table 2).

**Figure 5:**
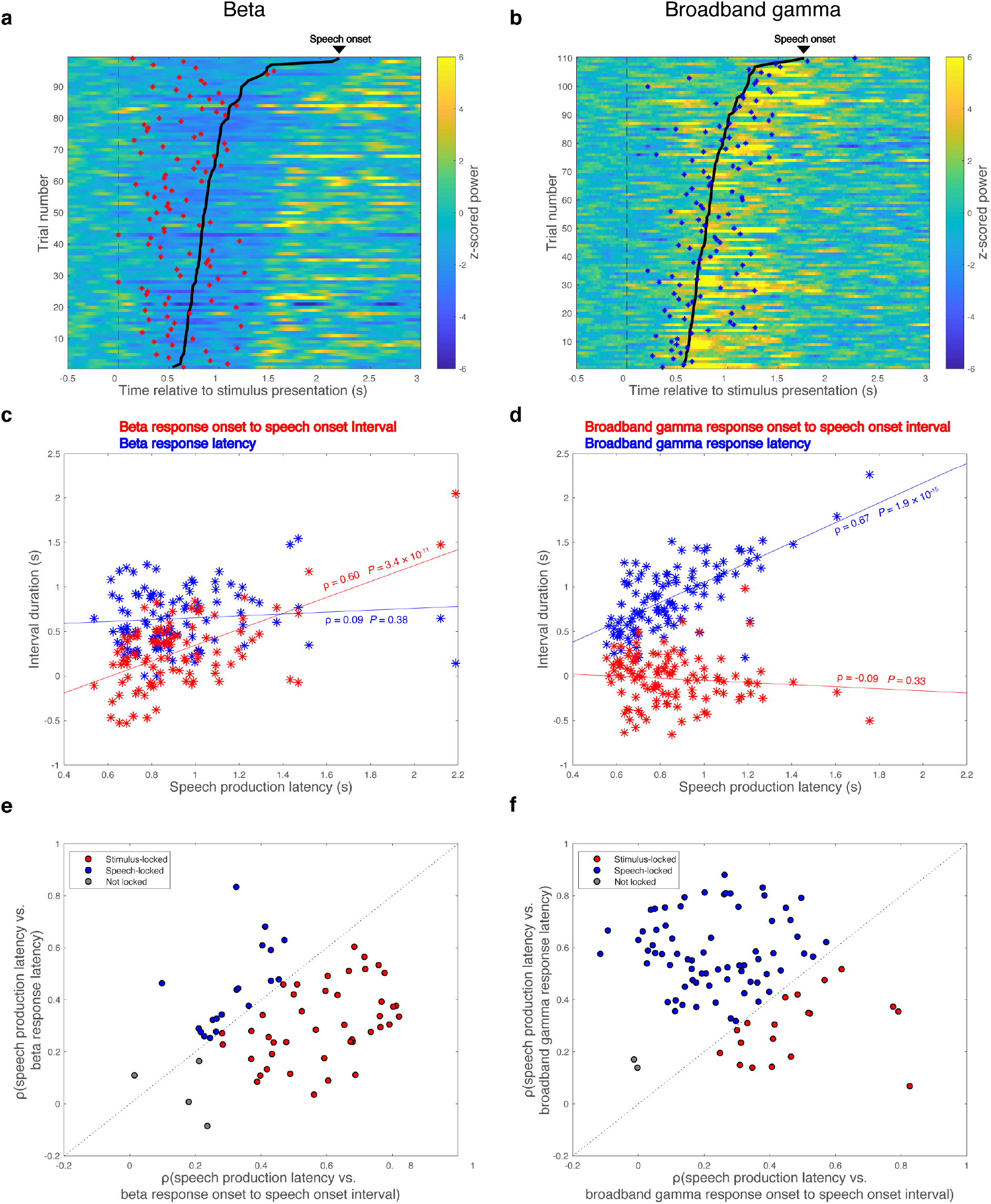
Beta decrease response is locked to stimulus presentation while broadband gamma increase response is locked to speech onset. **a**, **b** Raster plots for beta band responses (**a**) and broadband gamma responses (**b**) across trials of two representative recordings. Trials are aligned to stimulus presentation (indicated with black dashed lines) and sorted by speech production latency. Speech onsets are denoted with bold black lines. Onsets of significant beta activity decreases are marked with red asterisks in **a** and onsets of significant broadband gamma activity increases are marked with blue asterisks in **b**. **c**, **d** Band response onset to speech onset interval (red asterisks) and band response latency (blue asterisks) are correlated (Pearson’s correlation, α=0.05) with speech production latency respectively, for the two representative recordings. **e**, **f** The same correlation analysis is performed for all the recordings with significant beta decrease responses and all the recordings with significant broadband gamma increase responses, and the results are summarized in **e** and **f**, respectively. Recordings locked to stimulus presentation are shown in red, recordings locked to speech onset are shown in blue, and recordings not locked to either stimulus presentation or speech onset are shown in gray.

### Thalamic beta activity is selective to lexical status

To investigate the involvement and lateralization of the thalamus in lexical processing, only recordings that were from subjects with bilateral lead recordings (n=7) and that showed significant task-related modulation were included for lexicality-related analyses. As a result, 55 recordings (21 unilateral session left-side recordings, 34 from bilateral session left-side (20) and right-side (14) recordings) were included for beta lexical selectivity analysis. Nonword production was associated with a greater suppression of beta power compared with reading words. These differential beta responses were observed in both hemispheres (Figure 6a, c). In the left thalamus, significant word vs. nonword beta amplitude differences occurred between 0.8s and 1.8s after stimulus presentation (two-tailed paired t-test, n=41, *P*<0.05, Bonferroni corrected). Similarly, significant beta amplitude differences occurred in the right thalamus at 0.2-1.7s relative to stimulus presentation (two-tailed paired t-test, n=14, *P*<0.05, Bonferroni corrected).

**Figure 6:**
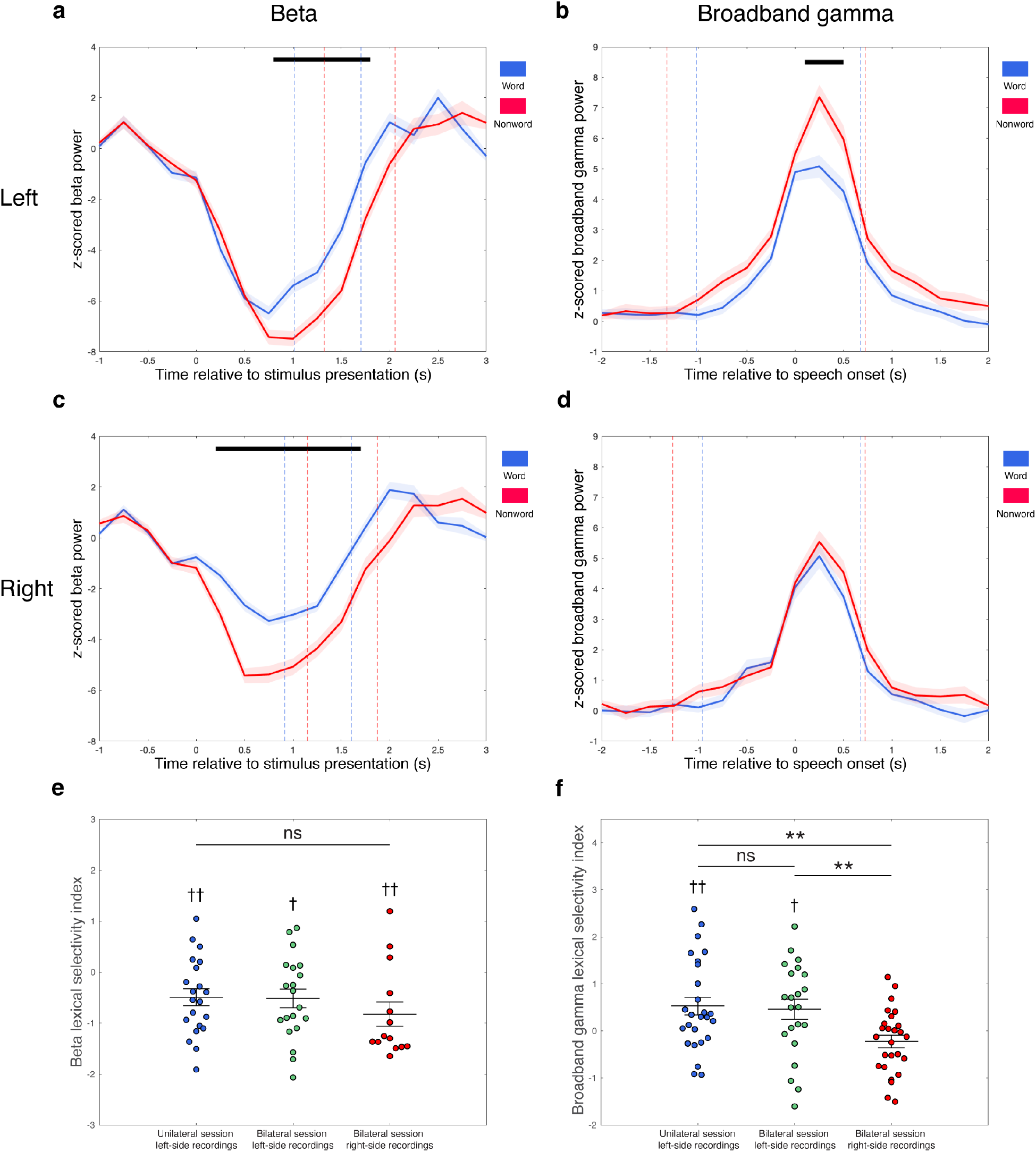
Differential band responses during word vs. nonword reading aloud in the left and right thalamus. **a**-**d** Comparisons of average band response amplitudes during word (blue) vs. nonword (red) reading aloud for beta band (**a**, **c**) and broadband gamma (**b**, **d**), in the left (**a**, **b**) and right thalamus (**c**, **d**). Band responses are averaged across trials of respective trial types and across recordings that showed significant task-related band responses in each side, aligned to stimulus presentation for beta band (**a**, **c**) and speech onset for broadband gamma (**b**, **d**). Average time points of speech onset and end of speech are marked with dashed lines in **a** and **c**, and average time points of stimulus presentation and end of speech are marked with dashed lines in **b** and **d**, for word trials (blue) and nonword trials (red) respectively. Black bars indicate significant differences of response amplitudes between word and nonword trials (two-tailed paired t-test, *P*<0.05, Bonferroni corrected). Standard errors are shaded in light colors. **e** Dot plot of beta lexical selectivity indexes of lead recordings that showed significant task-related beta decrease responses, grouped by recording sides and recording sessions (unilateral session left-side recordings in the first column, bilateral session left-side recordings in the second column, and bilateral session right-side recordings in the third column). Mean and SEM of beta lexical selectivity index across recordings of each group is superimposed on each column, respectively. **f** Dot plot of broadband gamma lexical selectivity indexes of lead recordings that had significant task-related broadband gamma increase responses, grouped by recording sides and recording sessions (unilateral session left-side recordings in the first column, bilateral session left-side recordings in the second column, and bilateral session right-side recordings in the third column). Mean and SEM of broadband gamma lexical selectivity index across recordings of each group is superimposed on each column, respectively. Twotailed one-sample t-test, †*P*<0.05, ††*P*<0.01; two-tailed two-sample t-test, \*\**P*<0.01, ns: not significant.

To quantify the lexical selectivity of thalamic beta activity, a lexical selectivity index was determined for each of these recordings. We utilized the time window of significant task-related beta activity decrease (−1.08-0.62s relative to speech onset (see Figure 4)) to calculate a mean task-related beta response power value for each trial, and then the values were compared between nonword trials vs. word trials in each recording by performing a two-sample t-test. The resulting t-statistic was the beta lexical selectivity index for this recording: a beta lexical selectivity index less than zero indicated stronger beta activity decrease (thus having more negative z-scored power) during nonword trials than during word trials, and vice versa. The mean beta lexical selectivity indexes for unilateral session left-side recordings (−0.49 ±0.76), bilateral session left-side (−0.52 ±0.82) and right-side (−0.82 ±0.89) recordings were all significantly less than zero (two-tailed one-sample t-test, t(20)=-2.97, *P*=0.0075; t(19)=-2.83, *P*=0.011; t(13)=-3.48, *P*=0.0041; Figure 6e), suggesting that the magnitude of the beta decrease was significantly nonword-selective in each case. There were no significant differences in beta lexical selectivity between recordings in the left and right thalamus, or between recordings from the unilateral and bilateral sessions on the left side (two-tailed two-sample t-test, unilateral session left-side recordings vs. bilateral session right-side recordings: t(33)=1.18, *P*=0.25; bilateral session left-side recordings vs. bilateral session right-side recordings: t(32)=1.05, *P*=0.30; unilateral session left-side recordings vs. bilateral session left-side recordings: t(39)=0.093, *P*=0.93).

### Lexical selectivity of thalamic broadband gamma activity is lateralized

74 recordings (26 unilateral session left-side recordings, 22 bilateral session left-side recordings, and 26 bilateral session right-side recordings) were included for broadband gamma lexical selectivity analysis. Significantly greater broadband gamma activity increases during nonword production were observed in the left thalamus, starting 0.1s after speech onset and persisting throughout the following 0.4s (two-tailed paired t-test, n=48, *P*<0.05, Bonferroni corrected; Figure 6b). In the right thalamus, however, the broadband gamma response curves observed during nonword and word reading were similar, without a significant difference in response amplitudes (two-tailed paired t-test, n=26, α=0.05, Bonferroni corrected; Figure 6d).

We next calculated broadband gamma lexical selectivity indexes, using the significant time window of −0.15-0.59s relative to speech onset (see Figure 4). A broadband gamma lexical selectivity index greater than zero indicated stronger broadband gamma activity increase during nonword trials than during word trials, and vice versa. The mean broadband gamma lexical selectivity indexes of both unilateral session left-side recordings (0.53 ±0.96) and bilateral session left-side recordings (0.46 ±1.00) were significantly greater than zero (two-tailed one-sample t-test, t(25)=2.80, *P*=0.0097; t(21)=2.14, *P*=0.044), indicating significant correlation of the magnitude of the gamma response to lexical status (Figure 6f). In contrast, broadband gamma responses in right side recordings did not show significant lexical selectivity (−0.22 ±0.68, two-tailed one-sample t-test, t(25)=-1.67, *P*=0.11). The differences in broadband gamma lexical selectivity between the left and right thalamus were demonstrated with two-tailed two-sample t-test (unilateral session left-side recordings vs. bilateral session right-side recordings: t(50)=3.25, *P*=0.0021; bilateral session left-side recordings vs. bilateral session right-side recordings: t(46)=2.79, *P*=0.0077), further suggesting that selectivity of thalamic broadband gamma activity to lexical status is lateralized to the left. Recordings in the unilateral and bilateral sessions on the left side did not differ in broadband gamma lexical selectivity (two-tailed two-sample t-test, t(46)=0.25, *P*=0.80), indicating the consistency of broadband gamma lexical selectivity between task sessions.

### Left side broadband gamma lexical selectivity is topographically organized

We observed that the recording sites in the left thalamus that were significantly nonword-selective in terms of broadband gamma response activity (broadband gamma lexical selectivity index>1.645 based on normal approximation of t-distribution) appeared to comprise more anterior locations in MNI space (Figure 7a). Pearson’s correlation tests demonstrated that the broadband gamma lexical selectivity index significantly correlated with recording location along the anterior-posterior axis (MNI-defined y coordinate; n=48, ρ=0.49, *P*=0.00041; Figure 7b) and the ventral-dorsal axis (MNI-defined z coordinate; n=48, ρ=0.30, *P*=0.041; Supplementary Figure 1a), but not the lateral-medial axis (MNI-defined x coordinate; n=48, ρ=-0.14, *P*=0.34; Supplementary Figure 1b) within the left thalamus. To avoid multicollinearity and to test out possible interactions between variables, a stepwise linear regression model was applied, which determined that the anterior-posterior location of the recording was the only significant predictor (estimated coefficient=0.17, SE=0.044, *P*=0.00041), while neither the ventral-dorsal location (*P*=0.72) nor the interaction between ventral-dorsal location and anterior-posterior location (*P*=0.14) explained the result. In order to control for subject differences and session differences, we further fitted linear mixed effects models to the data, entering subject and session as random effects. The results indicate that even after accounting for subject and session variability, the broadband gamma lexical selectivity index in the left thalamus had a significant gradient along the anterior-posterior axis, with greater broadband gamma lexical selectivity index more likely to be observed anteriorly (estimated coefficient=0.17, SE=0.043, *P*=0.00032). Based on the linear mixed effects modeling results, neither ventral-dorsal location (estimated coefficient=0.061, SE=0.035, *P*=0.087) nor lateral-medial location (estimated coefficient=-0.11, SE=0.093, *P*=0.24) of the recording had significant effect on broadband gamma lexical selectivity in the left thalamus. Taken together, these results suggest that broadband gamma lexical selectivity is dependent on the anterior-posterior location of the recording in the left ventral lateral thalamus, with greater nonword selectivity more likely to occur anteriorly.

**Figure 7:**
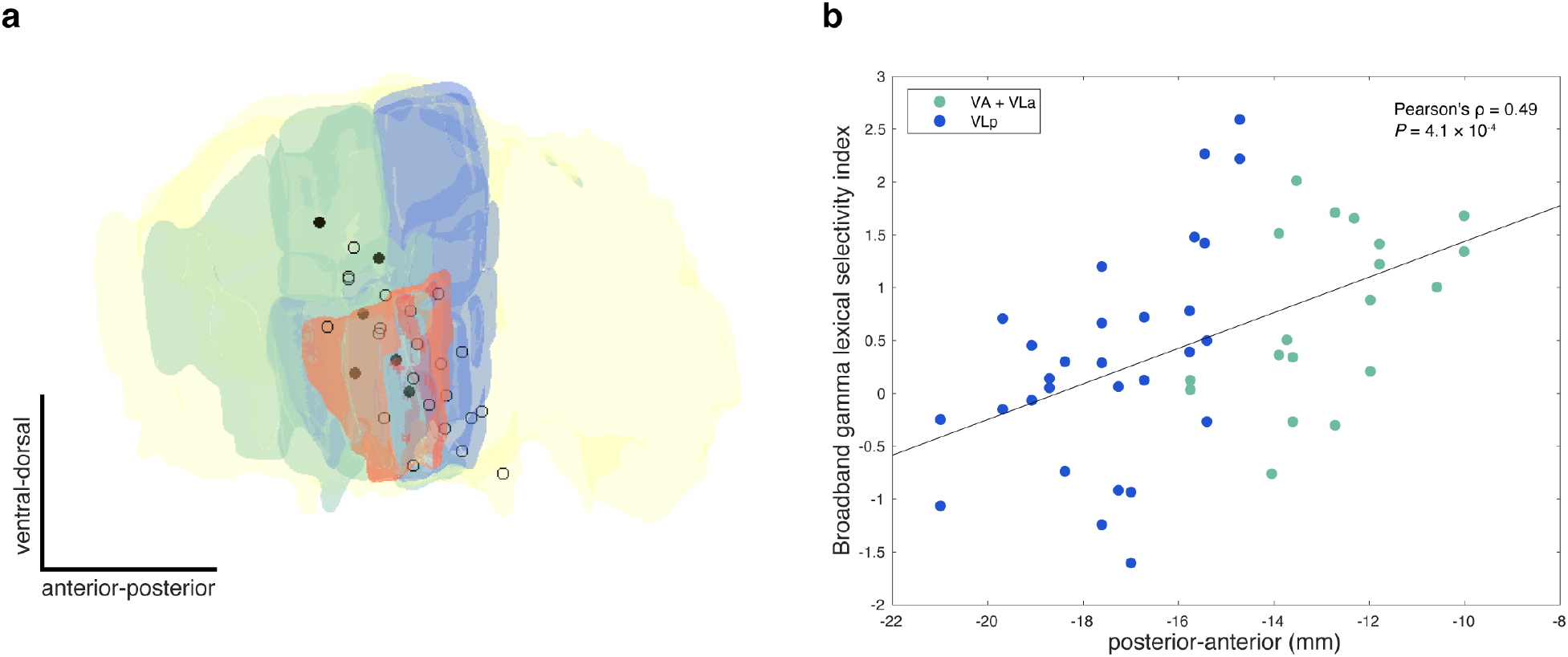
Broadband gamma lexical selectivity depends on anterior-posterior location of the recording in the left thalamus. **a** Left side lead recording sites are plotted together with anatomical structures (thalamus in yellow, VA and VLa in green, VLp in blue, and Vim in orange), viewed from a lateral direction. Recording sites where significantly nonword-selective recordings in terms of broadband gamma response amplitudes (broadband gamma lexical selectivity index>1.645 based on normal approximation of t-distribution) were observed in either session are filled with black, and remaining recording sites are shown in open circles. **b** Broadband gamma lexical selectivity indexes of left side lead recordings that showed significant task-related broadband gamma responses are correlated (Pearson’s correlation, α=0.05) with recording locations along the anterior-posterior axis (y coordinates in MNI space). Recordings inside VA or VLa are green-colored and recordings inside VLp are blue-colored.

No significant correlation (Pearson’s correlation, α=0.05) between broadband gamma lexical selectivity index and recording location was observed for right side recordings (Supplementary Figure 2 and 3), or for beta lexical selectivity on either side (Supplementary Figure 4 and 5), by any statistical modeling. These results suggest that the topography of lexical selectivity is unique to broadband gamma responses in the left thalamus.

## Discussion

Subcortical contributions to language production have been hypothesized largely from correlations of focal brain damage with aphasic syndromes and from language studies using functional magnetic resonance imaging (fMRI) or scalp electroencephalography (EEG) (Hebb & Ojemann, 2013). Our results are the first to demonstrate thalamic neural activity during reading aloud. We discovered that the encoding of lexical status is differentially represented in thalamic neural activity. Whereas greater beta activity decrease occurred during nonword trials as compared to word trials in both hemispheres, greater gamma increases in the left but not right thalamus were associated with the spoken production of nonwords as compared to words. Importantly, the level of broadband gamma lexical selectivity was greater in more anterior thalamic recording locations, regions more likely to receive basal ganglia than cerebellar afferents (Alexander et al., 1986).

We first observed that thalamic beta activity showed a task-related decrease, locked to stimulus presentation, consistent with the expected beta desynchronization that accompanies motor action. Beta oscillations are proposed to signal the maintenance of current sensorimotor and cognitive states (Engel & Fries, 2010), and event-related beta activity decreases during the preparation and the execution of voluntary movements have been observed across relevant brain regions, including the thalamus (Klostermann et al., 2007; Kühn et al., 2004; Paradiso, 2004; Tzagarakis et al., 2010). Beta activity was reduced across the entirety of each task event, which requires a series of underlying neural dynamics (processing the visual stimulus, retrieval of lexical information, encoding of the phonological information, and execution of articulatory movement), suggesting a state change that facilitated the process of speech production. Significant increases in broadband gamma activity began before but were predominantly locked to speech onset. Given that broadband gamma activity is thought to index synchronized local neuronal firing (Ray et al., 2008; Ray & Maunsell, 2011) and is associated with the functional activation of relevant brain regions during a wide range of sensorimotor and cognitive tasks, including speech and language processing (Bouchard et al., 2013; Chrabaszcz et al., 2019; Crone et al., 2006; Juphard et al., 2011; Uhlhaas et al., 2011), our data suggest that the ventral lateral thalamus tracks speech production.

Studies of the brain networks involved in lexical processing during reading primarily have been confined to cortical regions (Dietz et al., 2005; Fiez et al., 1999; Hagoort et al., 1999; Heim et al., 2005; Herbster et al., 1997; Juphard et al., 2011; Mechelli et al., 2003; Xu, 2001). In this study, we utilized the balanced design of words and nonwords in the stimulus sets and simultaneous bilateral recordings in the seven participants to determine whether lexical processing is differentially represented in the thalamus during reading aloud. Thalamic broadband gamma oscillations showed significantly stronger activation during nonword reading aloud than during word reading aloud, which evolved around speech onset and was present throughout the entire utterance. Notably, this broadband gamma lexical selectivity was found only in recordings from the left side. According to classic speech production models (Levelt et al., 1999), the time course of the observed broadband gamma activity difference between word and nonword trials corresponded to the articulatory stage of reading aloud. It is unlikely that this lexicality effect was motoric, considering that the motor complexity was balanced between word and nonword stimuli, and that this differential broadband gamma modulation was not equivalently observed in the right hemisphere. In fact, the dual-route theory of reading aloud has suggested that words and nonwords are read aloud differently: while real words can be read aloud via either grapheme-to-phoneme conversions or direct word-to-sound mapping, nonwords can only be read aloud via grapheme-to-phoneme conversions (Coltheart et al., 2001). It is possible that phonological encoding was still ongoing during nonword production, after a failed internal lexicon lookup procedure. Indeed, stronger gamma (50-150Hz) responses for pseudo-words than for words have been reported in Broca’s area (Brodmann’s areas 44 and 45) during silent word reading, where the length of this differential response increased with the length of the stimuli (Juphard et al., 2011). In addition, it is possible that reading aloud nonwords created a learning or error correction demand (Hickok, 2014). We propose that the lexicality effect on thalamic broadband gamma activity during reading may reflect left thalamic participation in phonological encoding, learning, and feedback monitoring during speech and language processing. Although previous studies have supported a lateralized thalamic role in language (Johnson & Ojemann, 2000), our results from simultaneous bilateral direct recordings in the thalamus are the first to provide direct supporting evidence.

A topography for broadband gamma lexical selectivity was observed in the left thalamus. The lexical selectivity of the broadband gamma response was significantly correlated with the anterior-posterior location of recording sites in the left thalamus, with a higher lexical selectivity more likely to appear anteriorly. In contrast, a gradient for broadband gamma lexical selectivity was not observed in the right thalamus. This finding further supports a unique language role lateralized to the left thalamus, and suggests functional heterogeneity in the left ventral lateral thalamus during speech and language processing. The anterior portion (VA and VLa) receives input primarily from the internal globus pallidus and substantia nigra pars reticulata, and has strong connections with frontal cortex, including Broca’s area (Alexander et al., 1986; Behrens et al., 2003; Bosch-Bouju et al., 2013; Hintzen et al., 2018; Hwang et al., 2017; Zhang et al., 2010). Broca’s area has consistently been associated with lexico-phonological processing (Dietz et al., 2005; Fiez et al., 1999; Hagoort et al., 1999; Heim et al., 2005; Herbster et al., 1997; Juphard et al., 2011; Mechelli et al., 2003; Xu, 2001). Thus, this thalamic region may participate in differential lexical processing during nonword vs. word reading aloud in concert with Broca’s area. In contrast, the posterior region (VLp), which receives cerebellar projections and preferentially sends output to primary motor cortex (Behrens et al., 2003; Bosch-Bouju et al., 2013; Hintzen et al., 2018; Hwang et al., 2017; Zhang et al., 2010), might be more related to motor control of a selected motor plan. This idea is supported by previous stimulation studies that have reported location-dependent effects of thalamic stimulation on speech and language: stimulation of VLa could cause acceleration of language processes, while stimulation of VLp often affected motor aspects of speech, such as perseveration and stuttering speech (Hebb & Ojemann, 2013). We note that VA has been proposed to participate in selection of a language unit during speech production, via basal ganglia-thalamo-cortical loop interactions (Crosson, 2013); it also is included in the “planning loop” in the GODIVA model of speech production (Bohland et al., 2010).

In contrast, although significantly stronger nonword-related than word-related beta activity decreases accompanied the reading aloud task bilaterally, neither lateralization nor topography was observed for thalamic beta lexical selectivity. Therefore, beta activity decreases likely represent nonspecific changes of cognitive and sensorimotor states that prepare the entire thalamo-cortical network for a behavioral change. Note that in the current work we did not try to compare task-related band response strength between different recording locations and different recording sides among subjects and sessions, because of a number of uncontrollable factors that affect the band oscillatory power (e.g. recording impedances and baseline neural activity are variable across subjects, sessions, and recording sides). In contrast, the lexical selectivity index, which was calculated by comparing word-related and nonword-related band response strength within each recording, is mostly independent of those factors and thus comparable at the group level. This idea is supported by the fact that lexical selectivity indexes in both beta and broadband gamma bands remained consistent across sessions.

In summary, our results are the first demonstration of time-frequency modulations of thalamic neural activity during reading aloud. These data suggest that lateralized and topographically organized thalamic pathways participate in speech production differently, based on whether a word or nonword is being read.

## Materials and methods

### Subjects

Eleven human subjects (3 females, 68.4 ±8.0 years) with essential tremor undergoing awake DBS implantation surgery targeting the Vim nucleus of the thalamus were studied. All participants were right-handed native English speakers. None had significant cognitive impairment based on a detailed neuropsychological evaluation performed during clinical evaluation for DBS surgery. All but one underwent bilateral (left side first) DBS lead implantation (one subject had one lead implanted in the left hemisphere previously and underwent right side lead implantation in the current study). Full demographic description of subjects is provided in Table 1. All protocols were approved by the Institutional Review Board of the University of Pittsburgh (IRB Protocol #PRO13110420), and all participants gave written informed consent.

**Table 1:** Subject characteristics.

| Subject | Age | Gender | Handedness | MMSE | LFP recording side | LFP recording type | Number of recordings | Mean number of trials per session | Trials rejected, % |
| --- | --- | --- | --- | --- | --- | --- | --- | --- | --- |
| 1 | 61 | Male | Right | NR | Left | Mapping electrodes | 12 | 111 | 7 |
| 2 | 70 | Female | Right | 30/30 | Right | Mapping electrodes | 6 | 107 | 11 |
| 3 | 66 | Male | Right | NR | Left | Mapping electrodes | 6 | 116 | 3 |
| 4 | 75 | Male | Right | 26/30 | Left | Mapping electrodes | 9 | 112 | 5 |
| 5 | 64 | Male | Right | 30/30 | Both | DBS leads | 12 | 104 | 13 |
| 6 | 53 | Male | Right | NR | Both | DBS leads | 12 | 113 | 5 |
| 7 | 67 | Male | Right | 26/30 | Both | DBS leads | 12 | 111 | 7 |
| 8 | 71 | Male | Right | 26/30 | Both | DBS leads | 12 | 108 | 9 |
| 9 | 84 | Female | Right | 30/30 | Both | DBS leads | 12 | 118 | 2 |
| 10 | 73 | Female | Right | 30/30 | Both | DBS leads | 12 | 109 | 9 |
| 11 | 68 | Male | Right | 28/30 | Both | DBS leads | 12 | 114 | 5 |

MMSE: Mini-Mental State Examination; NR: not recorded.

### Stimuli and experimental paradigm

Subjects performed a reading aloud task either in the course of subcortical mapping procedure during the surgery (4/11 subjects with mapping electrode recordings) or after the placement of DBS lead in each side (7/11 subjects with lead electrode recordings). For the 4 subjects with mapping electrode recordings, each subject performed up to 4 task sessions, while the 7 subjects with lead electrode recordings each performed 2 task sessions (the first session occurred after left lead was implanted, and the second session took place after bilateral implantation was completed). Each session included 120 trials. The stimuli were consonant-vowel-consonant (CVC) words and nonwords that were presented on a computer screen. Four lists of 120 stimuli (Supplementary Table 3) were constructed based on a previous work (Moore et al., 2017). The first 60 stimuli of each list alternated between unique words and nonwords, and those words and nonwords were balanced along a number of psycholinguistic features, including phoneme probability, phonological neighborhood density, bigram frequency, and biphone probability (see Supplementary Table 4 for the results of statistical comparisons (two-tailed two-sample t-test, n=60, α=0.05) between the two conditions in terms of these psycholinguistic parameters, which were calculated using the English Lexicon Project database (Balota et al., 2007)). The nonwords were duplicated twice to construct the last 60 stimuli of each list. One of the four stimulus lists was presented to the subjects during each task session. For lexicality-related analyses, only the first 60 trials of each session were used.

The experimental paradigm was programmed using MATLAB (MathWorks, Natick, MA) and Psychophysics Toolbox extensions (Brainard, 1997). A schematic of the experiment is shown in Figure 1. Before each trial, a white fixation cross was presented on the screen. Each trial was initiated manually by the experimenter, with the appearance of a green fixation cross on the screen. The green fixation cross lasted 250ms, and was followed by a variable time delay (500-1000ms) during which the screen remained black. Then the CVC syllable stimulus was presented in white on the screen, and subjects were instructed to read it aloud. The text remained on the screen until the subjects finished speaking. A white fixation cross was presented on the screen during the inter-trial interval (ITI).

### Electrophysiological recordings

For the 4/11 subjects where physiological subcortical mapping was administered, LFP recordings were performed using the Neuro-Omega recording system (Alpha Omega, Nazareth, Israel) and mapping electrodes that have a stainless steel macroelectrode ring (0.55mm in diameter, 1.4mm in length) 3mm above the tip, while in the other 7/11 subjects LFP signal was recorded from Medtronic Model 3387 DBS leads (Medtronic, Minneapolis, MN) with four platinum-iridium electrodes (1.27mm in diameter, 1.5mm in length) that are spaced 1.5mm apart, using Grapevine Neural Interface Processor (Ripple LLC, Salt Lake City, UT). The mapping electrodes and the DBS leads targeted the Vim. For subjects with mapping electrode recordings, three mapping electrodes were placed in three trajectories (anterior, central, and posterior or central, posterior, and medial) of a standard cross-shaped Ben-Gun array with a 2mm center-to-center spacing, and made simultaneous recordings starting at 15mm above the surgical target with manual advance in 0.1mm steps. The reading aloud task was carried out in pauses during subcortical mapping procedure and subjects performed in up to four recording sessions, with each session corresponding to a different recording depth. Mapping electrode recordings were performed only during the left side implantation except for one subject who underwent unilateral implantation in the right side, and thus got mapping electrode recordings only in the right side. For subjects with DBS lead electrode recordings, the task was administered in two recording sessions, one after the implantation of the DBS lead in the left side, receiving recordings from only the left DBS lead electrodes, and the other after bilateral DBS lead implantation was completed, receiving simultaneous recordings from bilateral lead electrodes. The LFP signal recorded from mapping electrodes was sampled at 44kHz and bandpass filtered from 0.075Hz to 10kHz, and data recorded from DBS lead electrodes were collected at 30kHz. Signal collected in one recording site in one session counted as one recording. Subject recording characteristics are summarized in Table 1.

### Audio recordings

Subjects’ speech signal was recorded using an omnidirectional microphone (Audio-Technica model ATR3350iS Mic, frequency response 50-18,000Hz (Audio-Technica, Machida, Japan) for 6 subjects, and PreSonus model PRM1 Precision Flat Frequency Mic, frequency response 20-20,000Hz (PreSonus, Baton Rouge, LA) for 5 subjects) placed approximately 8cm away from the subject’s mouth and oriented at an angle of about 45°. The audio signal was collected by Grapevine Neural Interface Processor at a sampling frequency of 30kHz. For subjects with mapping electrode recordings, the audio signal was then synchronized with neural signal recorded by the Neuro-Omega system using digital pulses delivered to both recording systems via a USB data acquisition unit (model USB-1208FS, Measurement Computing, Norton, MA).

### Electrode localization

DBS lead electrodes and mapping electrodes were localized using LEAD-DBS toolbox (Horn et al., 2019; Horn & Kühn, 2015). Postoperative brain scans were coregistered to preoperative brain scans using open-source Advanced Normalization Tools (ANTs). Pre- and postoperative acquisitions were then normalized into MNI ICBM152 NLIN 2009b stereotactic space (Fonov et al., 2011). Both coregistration and normalization results underwent manual quality check. Semi-automatic reconstruction of electrodes in MNI space was performed in LEAD-DBS and MNI-defined coordinates were determined for all the electrode contacts (Figure 2). A digitized and normalized to MNI space version of the Ewert atlas (Ewert et al., 2018) was used to categorize the electrode contacts. A contact was assigned to a nucleus if it was within or in the vicinity of the nucleus (1mm cut-off) based on the minimum Euclidean distance between the contact and the voxels of the nucleus.

### Data pre-processing

The audio signal was segmented into trials and the speech sound was coded by communication science students trained in phonetics in a custom-designed graphical user interface implemented in MATLAB. The coding results were manually checked by a speech-language pathologist. For each trial, (1) the onset of speech was identified, (2) the end of speech was identified, and (3) the speech content was identified. Trials were considered to have correct speech responses and were included in further analyses if they met all the following criteria: (1) the subject’s speech response could be clearly identified by the coder, (2) the subject’s response was a CVC syllable consisting of the targeted phonemes, and (3) the response did not make nonword into a word or word into a nonword.

Electrophysiological data were pre-processed using custom code based on FieldTrip toolbox (Oostenveld et al., 2011) in MATLAB. The data were resampled at 1kHz and band-pass filtered from 2 to 400Hz. The data were also notch-filtered at 60Hz and its’ harmonics to remove line noise. Time series data from all recording sites were visually and quantitatively inspected for quality control. The data were then segmented into trials, each spanning 2s before stimulus presentation to 2s after the end of speech. Trials with artefacts or excessive noise were identified both manually and quantitatively, and were excluded for subsequent analyses. Combined with trials that did not meet the criteria for correct speech responses, an average 6.9 ±3.4% of trials per subject were rejected (Table 1). The remaining data were common-average referenced to minimize noise. For spectral-temporal analysis, the data were decomposed using Morlet wavelet transformation (width=7) over frequencies of 2Hz to 200Hz in increment steps of 2Hz. For band activity analyses, instantaneous analytic amplitudes of beta and broadband gamma frequency bands were extracted from respective bandpass filters using Hilbert transform (MATLAB function *hilbert*). The resulting signal of each trial was z-scored relative to the baseline (a period of 1000ms preceding stimulus presentation).

### Task-related responses

Time-frequency data were averaged across all trials centering on speech onset (from 2s before speech onset to 2s after speech onset), z-scored to baseline, and then averaged across all recordings. Nonparametric two-tailed Wilcoxon signed-rank test was performed to determine significant time-frequency modulations compared to baseline (n=117, α=0.05, Bonferroni corrected). Significant time windows for beta and broadband gamma determined by the statistical results were used to calculate a mean beta response strength and mean broadband gamma response strength for each trial in each recording. Then one-tailed one-sample t-test was performed on each recording to determine recordings that had significant beta activity decreases and recordings that had significant broadband gamma activity increases, respectively (α=0.05, Bonferroni corrected).

### Locking analysis

In an effort to characterize the timing properties of beta decrease response and broadband gamma increase response for recordings with significant task-related changes in either band activity, we examined the trial-to-trial relationships of significant band response onsets to stimulus presentation versus speech onset in these recordings. First, beta or broadband gamma time series data of each trial were smoothed using a moving average kernel of 200ms (MATLAB function *smoothdata*) and z-scored to baseline, in order to minimize single-trial noise. For recordings with significant beta decrease responses, a thresholding method with a critical value of z=-1.645 was applied to determine the onset of beta response for each trial. Specifically, for each trial, the period with band response power below the threshold that gave minimum summed activity was considered as activation period and the beginning of the period was determined as onset of beta response. For recordings with significant broadband gamma increase responses, the onset of broadband gamma increase was determined in a similar way, except that a critical value of z=1.645 was used and period of maximum summed activity above the threshold was considered as activation period. For each band, a trial of one recording was discarded for locking analysis if no beyond-threshold period was present throughout the trial. Two intervals were calculated for each trial: (1) the interval between stimulus presentation and the onset of significant band response (band response latency), and (2) the interval between the onset of significant band response and the onset of speech. Then the two intervals were correlated (Pearson’s correlation, α=0.05) with speech production latency (interval between stimulus presentation and onset of speech) across trials for each recording, respectively. The band response of a recording was considered to be more time-locked to stimulus presentation, if (1) band response onset to speech onset interval was significantly correlated with speech production latency, and (2) the correlation coefficient (Pearson’s ρ) between band response onset to speech onset interval and speech production latency was greater than the correlation coefficient between band response latency and speech production latency. On the contrary, the band response was considered to be more time-locked to speech onset, if (1) band response latency was significantly correlated with speech production latency, and (2) the correlation coefficient between band response latency and speech production latency was greater than the correlation coefficient between band response onset to speech onset interval and speech production latency. If a band response did not meet any of the two criteria, it was considered not locked to either stimulus presentation or speech onset. χ^2^ tests were performed to differentiate the locking properties of broadband gamma increase response and beta decrease response, and to test if the locking properties differed between recording sides (α=0.05).

### Lexical selectivity analysis

We utilized the first 60 trials of recordings that were from the seven subjects with bilateral data and that showed significant task-related modulations in beta or broadband gamma activity to make comparisons between the left and right thalamus in terms of band lexical selectivity. A mean z-scored band response curve averaged across trials and across recordings was obtained for each frequency band, each trial type (word/nonword), in each side. Trials were aligned to stimulus presentation for beta and aligned to speech onset for broadband gamma, based on their respective locking properties. Periods of significant differences between word-related responses and nonword-related responses were determined with two-tailed paired t-test using a sliding window of 100ms (α=0.05, Bonferroni corrected).

To quantify the extent of band response difference between word and nonword trials, beta lexical selectivity indexes and broadband gamma lexical selectivity indexes were calculated for respective recordings. For each recording, the mean response power values of a particular frequency band over the corresponding significant time window determined before (see **Task-related responses**) were calculated for word trials and nonword trials. Two-sample t-test was then performed between nonword-related power values and word-related power values, and the resulting t-statistic was the lexical selectivity index of that frequency band for that recording. For beta, a lexical selectivity index smaller than zero meant that the recording showed stronger beta activity decrease (thus having more negative z-scored power) during nonword trials than during word trials, and vice versa. For broadband gamma, a lexical selectivity index greater than zero indicated that the recording showed stronger broadband gamma activity increase during nonword reading aloud than during word reading aloud, and vice versa. The beta or broadband gamma activity of a recording was considered significantly nonword-selective, if lexical selectivity index was smaller than −1.645 for beta or greater than 1.645 for broadband gamma (normal approximation of t-distribution). Two-tailed one-sample t-test was performed to test if lexical selectivity indexes of a selected group of recordings were significantly different from zero (α=0.05). Two-tailed two-sample t-test was performed to test if lexical selectivity indexes significantly differed between two groups of recordings (α=0.05).

### Analysis of location dependency of lexical selectivity

We further sought to examine possible dependence of band lexical selectivity on recording location, for both beta band and broadband gamma and in both left and right recording sides. Several regression models were applied to the data. First, simple linear regression (Pearson’s correlation, α=0.05) was performed to correlate lexical selectivity index with recording location: anterior-posterior location (MNI-defined y coordinate), ventral-dorsal location (MNI-defined z coordinate), and lateral-medial location (MNI-defined x coordinate), respectively. A series of stepwise linear regression models (MATLAB function *stepwiselm*, response variable: lexical selectivity index, predictor variables: MNI-defined x, y, z coordinates and their interactions, α=0.05) were then carried out to test out possible variable interactions, minimize multicollinearity, and determine the final location dependency model of band lexical selectivity. Finally, in order to account for possible lexical selectivity differences that might exist due to subject- and session-specific variations, linear mixed effects models (MATLAB function *fitglme*) were also applied to the data, setting subject and session as random effects.

## Acknowledgments

This work was supported by NIH grant U01NS098969 to R.M.R., the Hamot Health foundation to R.M.R., and a University of Pittsburgh Brain Institute NeuroDiscovery Pilot Research Award to R.M.R. This work was supported by the University of Pittsburgh-Tsinghua University Scholars Program to D.W.

## Supplementary Information

**Supplementary Figure 1:**
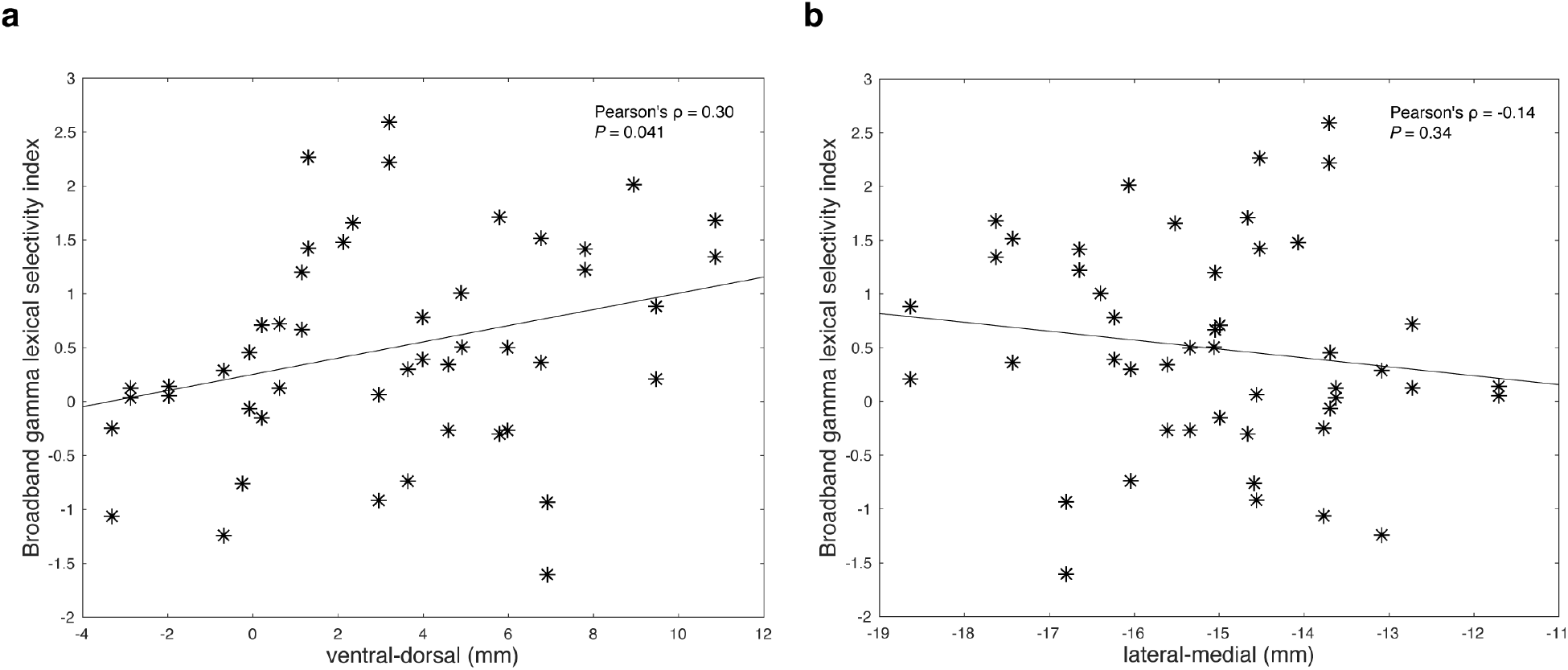
Correlation between left thalamic broadband gamma lexical selectivity and recording locations along ventral-dorsal and lateral-medial axes. Broadband gamma lexical selectivity indexes of left side lead recordings that showed significant task-related broadband gamma increase responses are correlated (Pearson’s correlation, α=0.05) with ventral-dorsal locations (**a**) and lateral-medial locations (**b**) of the recordings, respectively.

**Supplementary Figure 2:**
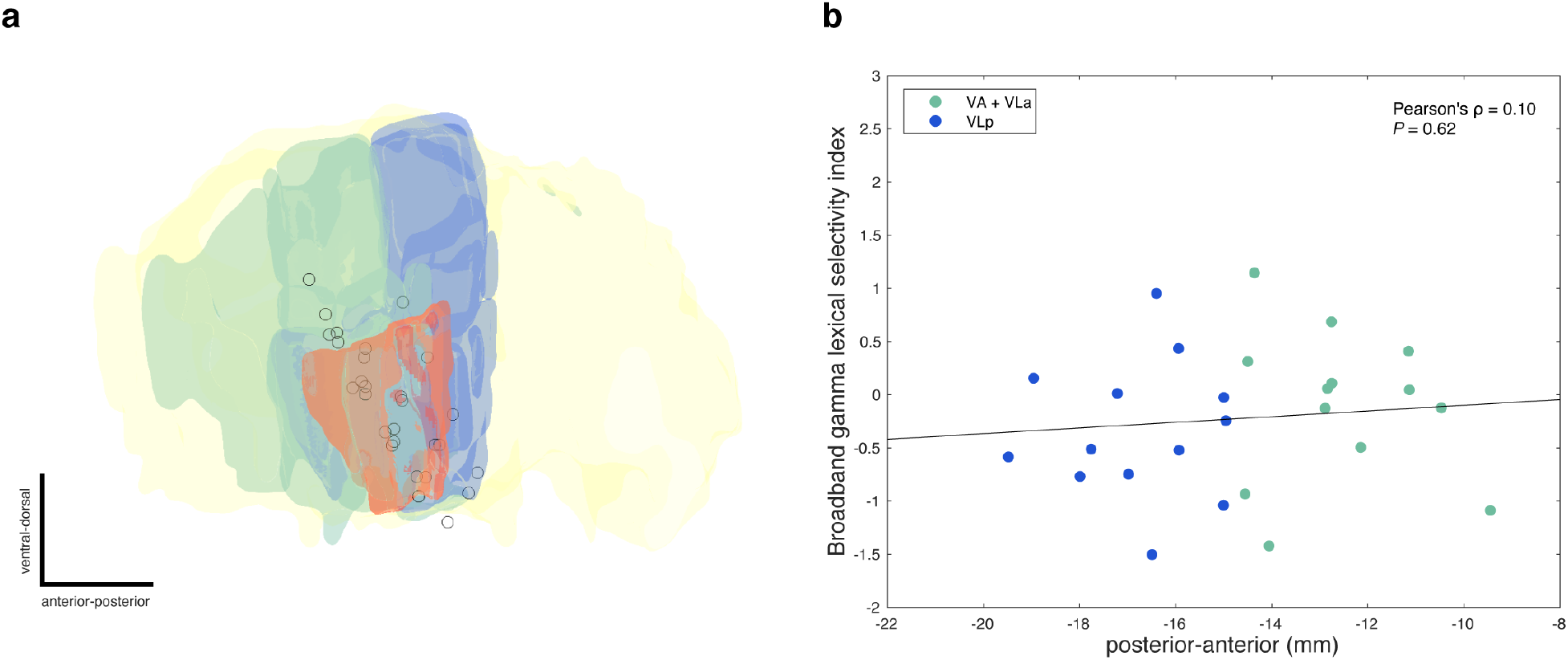
Broadband gamma lexical selectivity does not depend on anterior-posterior location of the recording in the right thalamus. **a** Right side lead recording sites are flipped to the left and plotted together with anatomical structures (thalamus in yellow, VA and VLa in green, VLp in blue, and Vim in orange), viewed from a lateral direction. Recording sites where significantly nonword-selective recordings in terms of broadband gamma response amplitudes (broadband gamma lexical selectivity index>1.645 based on normal approximation of t-distribution) were observed are filled with black (n=0), and remaining recording sites are shown in open circles. **b** Broadband gamma lexical selectivity indexes of right side lead recordings that showed significant task-related broadband gamma increase responses are correlated (Pearson’s correlation, α=0.05) with recording locations along the anterior-posterior axis. Recordings inside VA or VLa are green-colored and recordings inside VLp are blue-colored.

**Supplementary Figure 3:**
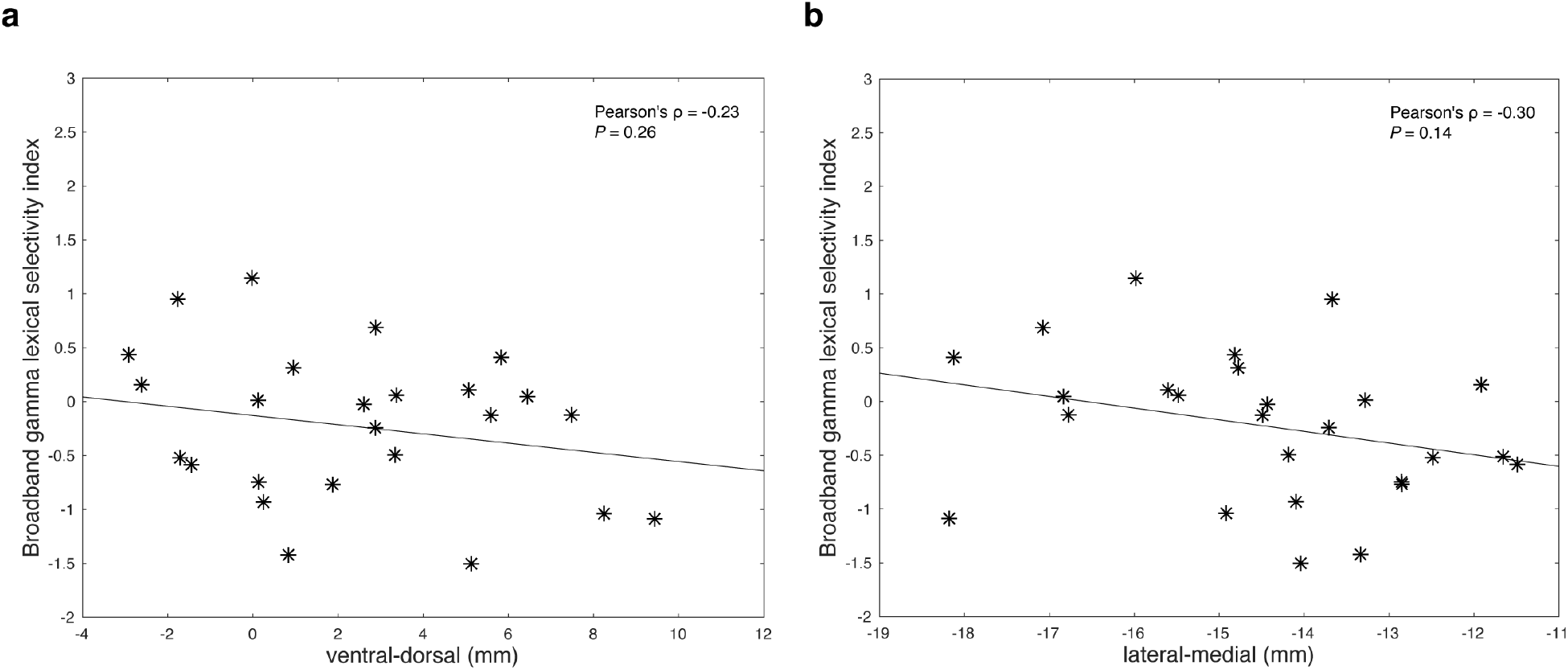
Correlation between right thalamic broadband gamma lexical selectivity and recording locations along ventral-dorsal and lateral-medial axes. Broadband gamma lexical selectivity indexes of right side lead recordings that showed significant task-related broadband gamma increase responses are correlated (Pearson’s correlation, α=0.05) with ventral-dorsal locations (**a**) and lateral-medial locations (**b**) of the recordings, respectively.

**Supplementary Figure 4:**
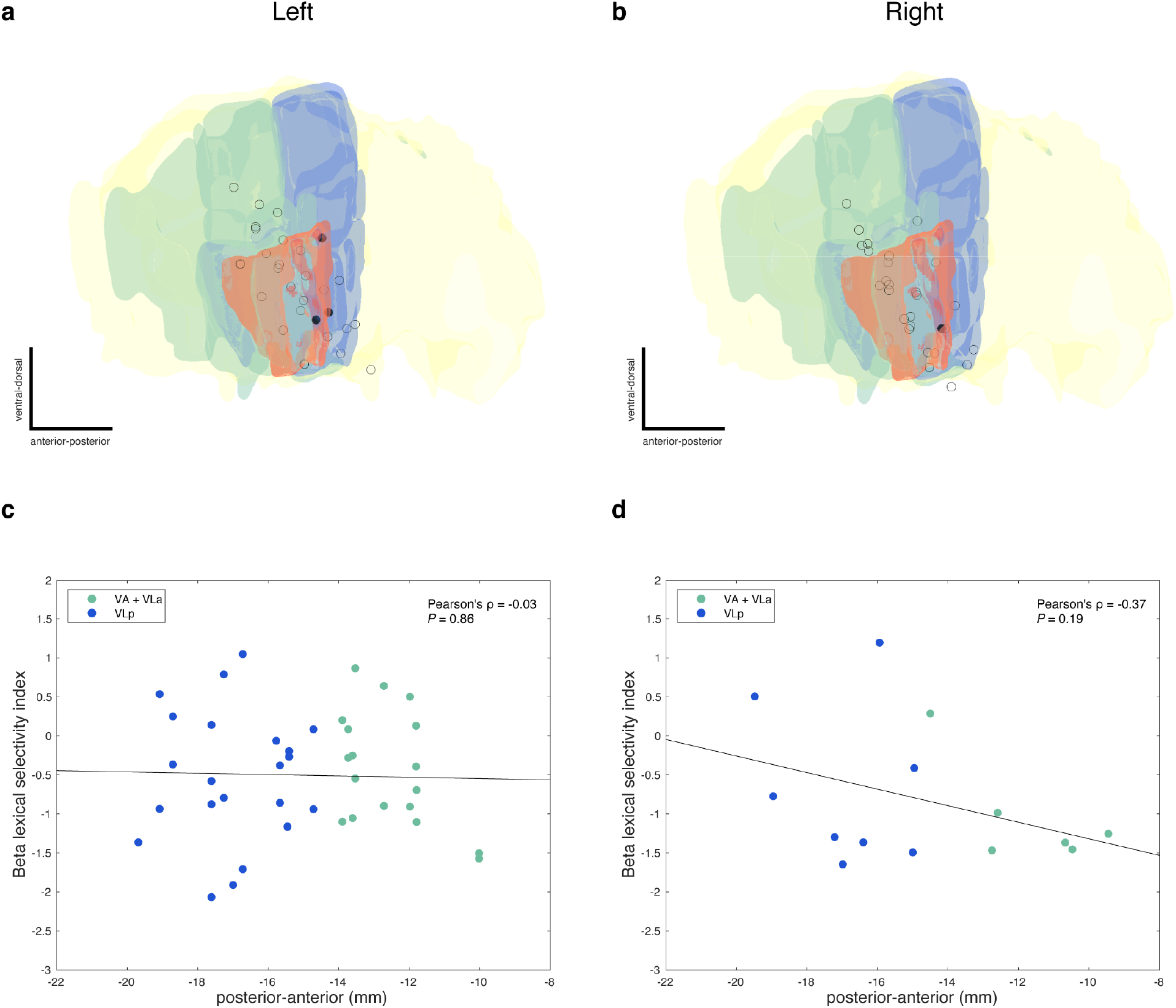
Beta lexical selectivity does not depend on anterior-posterior location of the recording in either side. **a** Left side lead recording sites are plotted together with anatomical structures (thalamus in yellow, VA and VLa in green, VLp in blue, and Vim in orange), viewed from a lateral direction. Recording sites where significantly nonword-selective recordings in terms of beta response amplitudes (beta lexical selectivity index<-1.645 based on normal approximation of t-distribution) were observed in either session are filled with black, and remaining recording sites are shown in open circles. **b** Right side lead recording sites are flipped to the left and plotted in the same way. **c** Beta lexical selectivity indexes of left side lead recordings that showed significant task-related beta decrease responses are correlated (Pearson’s correlation, α=0.05) with recording locations along the anterior-posterior axis. **d** Beta lexical selectivity indexes of right side lead recordings that showed significant task-related beta decrease responses are correlated (Pearson’s correlation, α=0.05) with recording locations along the anterior-posterior axis. In **c** and **d**, recordings inside VA or VLa are green-colored and recordings inside VLp are blue-colored.

**Supplementary Figure 5:**
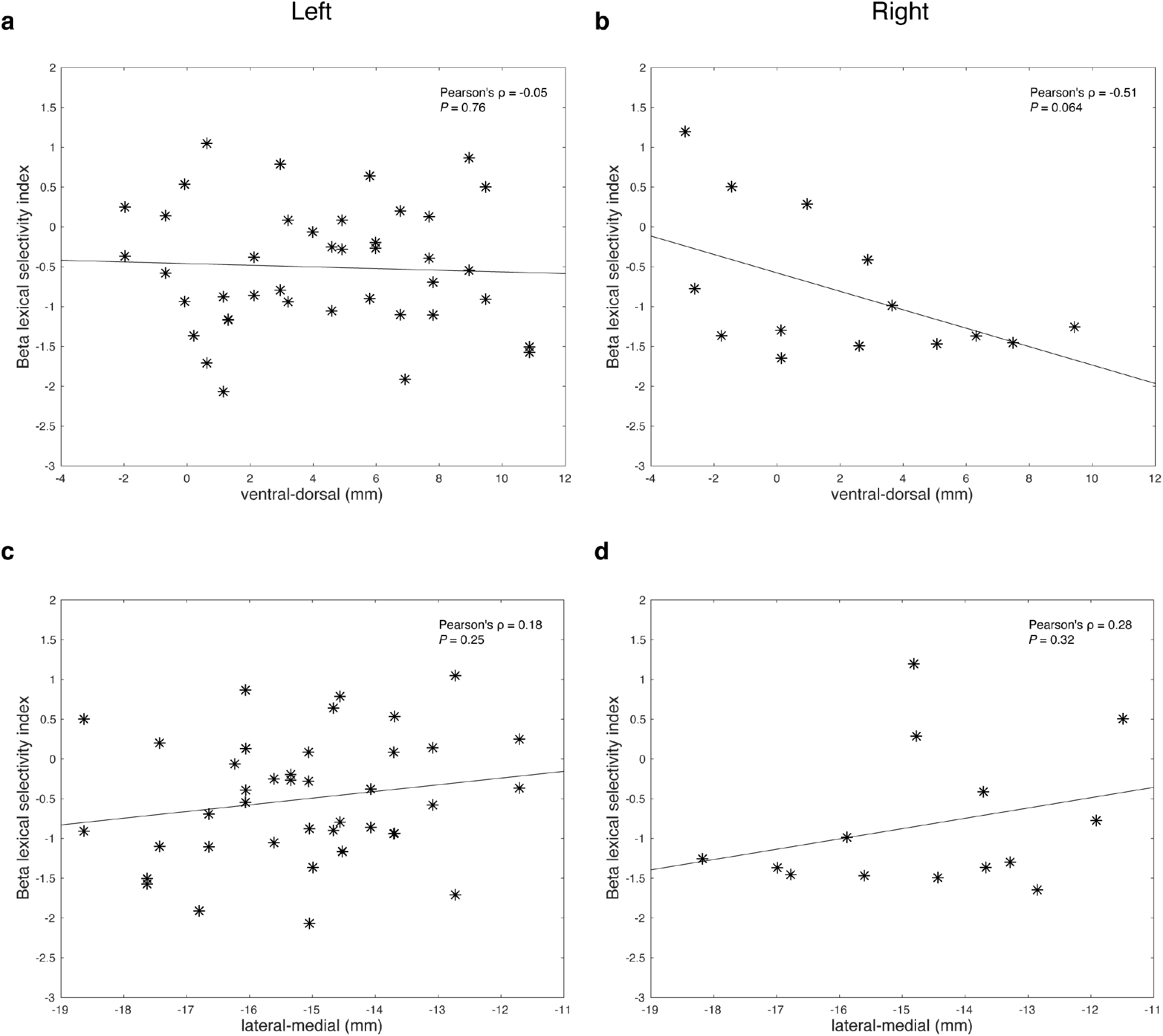
Correlation between thalamic beta lexical selectivity and recording locations along ventral-dorsal and lateral-medial axes. **a**, **c** Beta lexical selectivity indexes of left side lead recordings that showed significant task-related beta decrease responses are correlated (Pearson’s correlation, α=0.05) with ventral-dorsal locations (**a**) and lateral-medial locations (**c**) of the recordings, respectively. **b**, **d** Beta lexical selectivity indexes of right side lead recordings that showed significant task-related beta decrease responses are correlated (Pearson’s correlation, α=0.05) with ventral-dorsal locations (**b**) and lateral-medial locations (**d**) of the recordings, respectively.

**Supplementary Table 1:** Broadband gamma increase response and beta decrease response differ in locking properties.

| Response type | Number of recordings | Stimulus-locked (%) | Speech-locked (%) | Not locked (%) |
| --- | --- | --- | --- | --- |
| Broadband gamma increase response | 91 | 19 (21) | 70 (77) | 2 (2) |
| Beta decrease response | 66 | 43 (65) | 19 (29) | 4 (6) |
| $\chi^2$ | - | 31.4 | 36.1 | 1.55 |
| $P$ | - | $< 10^{-5}$ | $< 10^{-5}$ | 0.21 |

**Supplementary Table 2:** Locking properties of broadband gamma increase response and beta decrease response do not differ between recording sides.

| Response type | Number of recordings | Stimulus-locked (%) | Speech-locked (%) | Not locked (%) |
| --- | --- | --- | --- | --- |
| Broadband gamma increase response |  |  |  |  |
| Left | 59 | 10 (17) | 47 (80) | 2 (3) |
| Right | 32 | 9 (28) | 23 (72) | 0 (0) |
| $\chi^2$ | 2.49 | | | |
| $P$ | 0.29 | | | |
| Beta decrease response |  |  |  |  |
| Left | 49 | 31 (63) | 14 (29) | 4 (8) |
| Right | 17 | 12 (71) | 5 (29) | 0 (0) |
| $\chi^2$ | 1.50 | | | |
| $P$ | 0.47 | | | |

**Supplementary Table 3:**
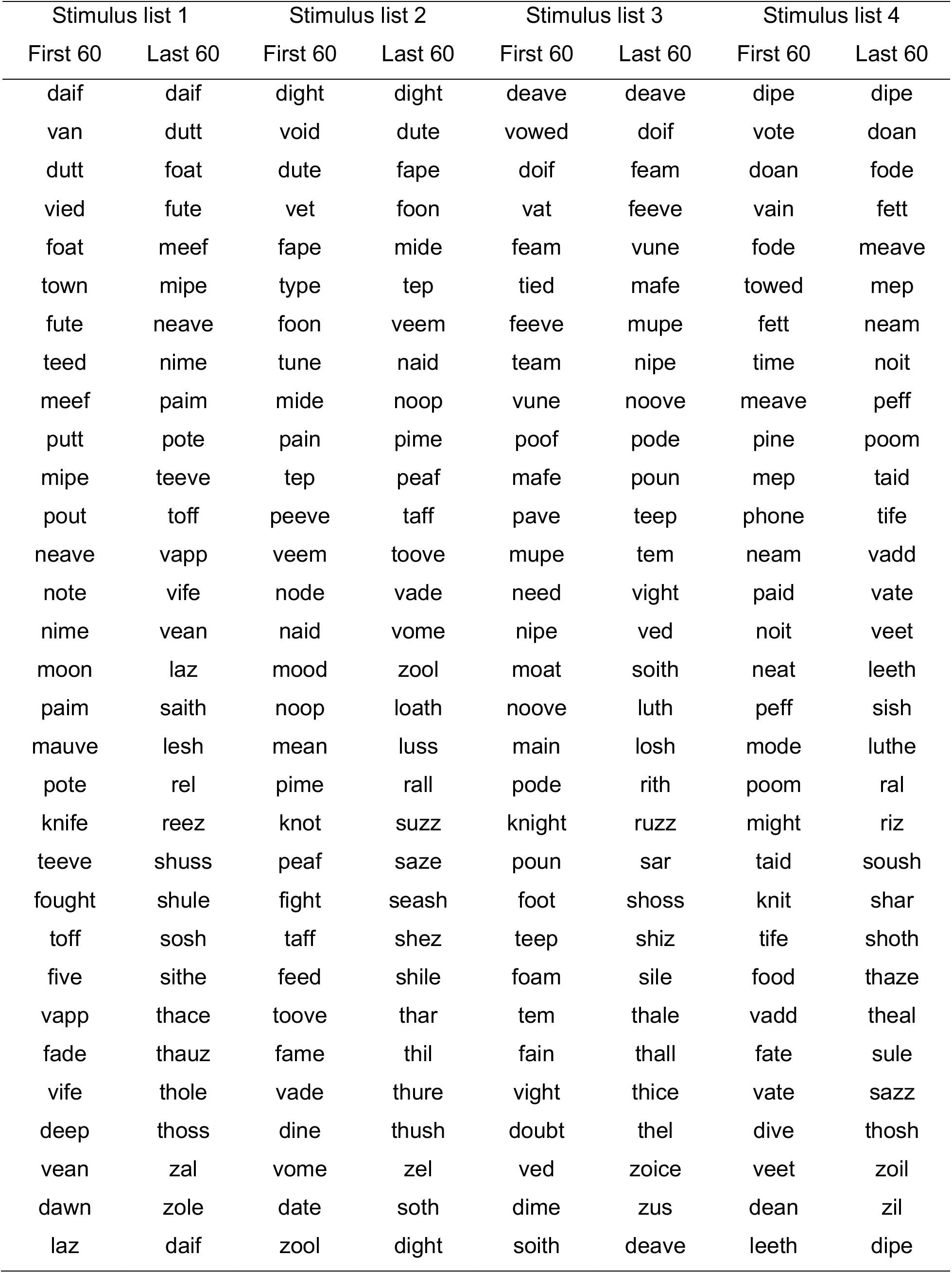
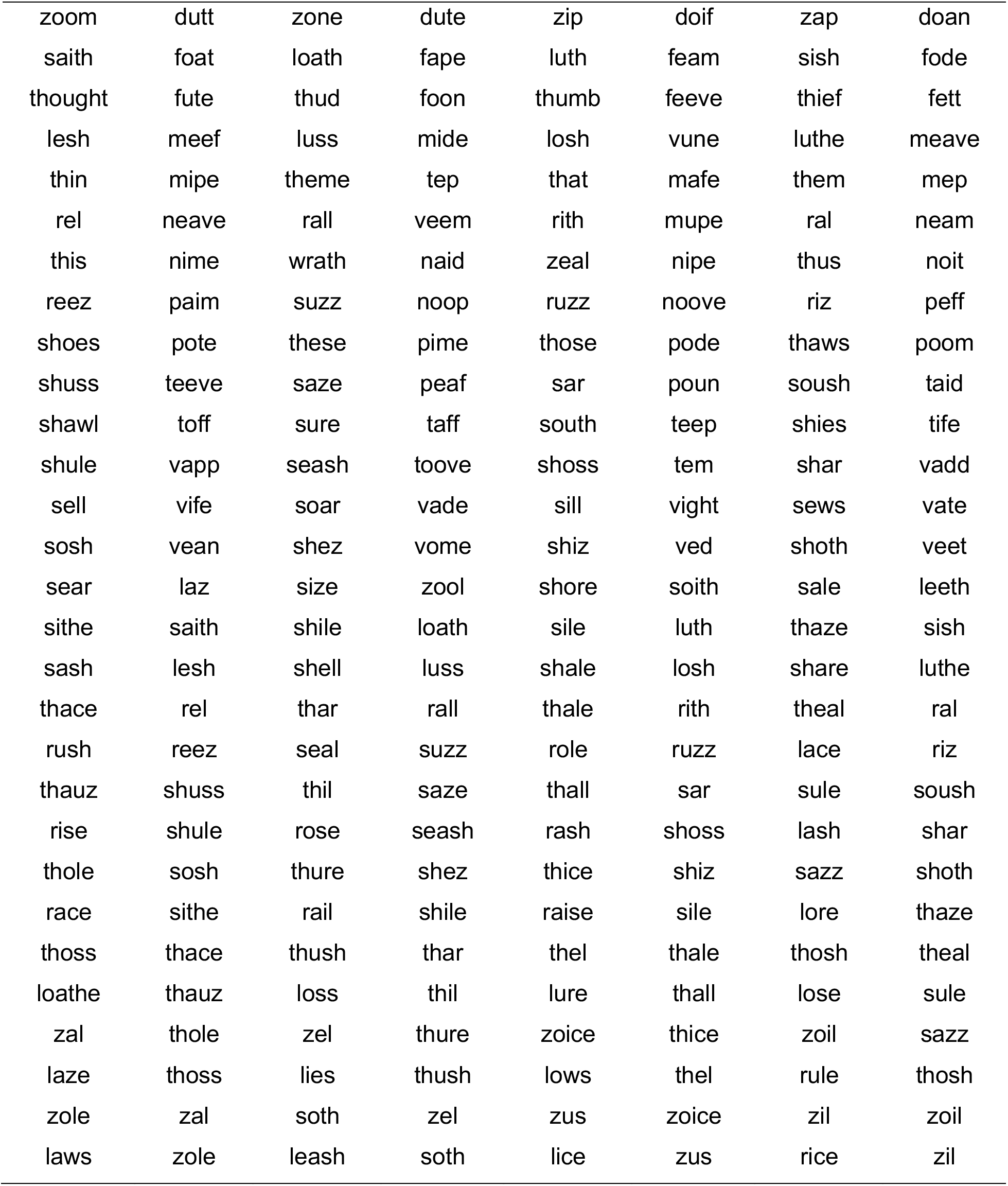
Stimulus lists.

**Supplementary Table 4:**
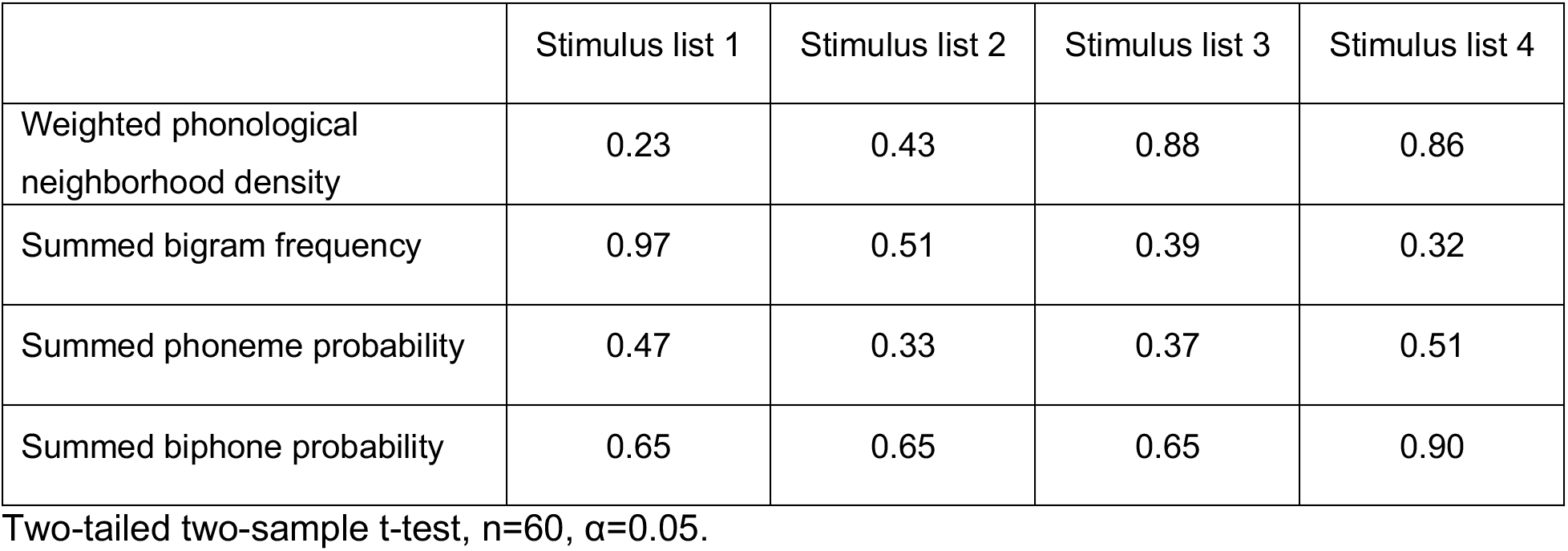
*P* values of statistical tests comparing psycholinguistic parameters between word and nonword stimuli.

